# iSubGen: Integrative Subtype Generation by Pairwise Similarity Assessment

**DOI:** 10.1101/2021.05.13.444087

**Authors:** Natalie S. Fox, Syed Haider, Constance H. Li, Paul C. Boutros

## Abstract

There are myriad types of biomedical data– genetics, transcriptomics, clinical, imaging, wearable devices and many more. When a group of patients with the same underlying disease exhibit similarities across multiple types of data, this is called a subtype. Disease subtypes can reflect etiology and sometimes predict clinical behaviour. Existing subtyping approaches struggle to simultaneously handle multiple diverse data types, particularly when there is missing information, as is common in most real-world clinical datasets. To improve subtype discovery, we exploited changes in the correlation-structure between different data types to create iSubGen, an algorithm for <u>i</u>ntegrative <u>sub</u>type <u>gen</u>eration. iSubGen can combine arbitrary data types for subtype discovery, such as merging molecular, mutational signature, pathway and micro-environmental data. iSubGen recapitulates known subtypes across multiple diseases, even in the face of substantial missing data. It identifies groups of patients with divergent clinical outcomes, and can combine arbitrary data types for subtype discovery, such as merging molecular, mutational signature, pathway and micro-environmental data. iSubGen can accommodate any feature that can be compared with a similarity-metric, and provides a versatile approach for creating subtypes. It is available at https://CRAN.R-project.org/package=iSubGen.

## Introduction

Most diseases show substantial inter-patient variability in presentation, progression and response to treatment; this heterogeneity is a hallmark of cancer, autoimmune disorders and neurological disorders, amongst others^1–6^. These differences often reflect common patterns of disease features, called subtypes, which can be important for clinical management by reducing the heterogeneity in presentation and progression^2,7,8^.

Disease subtypes play a particularly important role in cancer, where almost all tumours arise from a single cell, and features of that cell shape tumour initiation, progression and evolution^9,10^. The location of the primary cancer lesion influences the types of interventions possible and their efficacies, leading cancers to be grouped clinically based on their tissue of origin. Individual tissues contain cells of different types and distinct gene expression landscapes, and these evolve into cancers with distinct characteristics^11^. Further, cells of a single cell-type can lead to different types of cancer based on the identity and timing of driver mutations, and on the microenvironmental pressures they experience during tumourgenesis^10,12,13^.

These variable, but repeatedly observed, evolutionary courses of cancers originating in a single anatomical location are termed “cancer subtypes”. Historically, cancer subtypes have been defined histopathologically^3–6^. More recently, high-throughput molecular assays have discovered and defined subtypes^7,14–17^. Both approaches can identify groups of cancers with less heterogeneous prognoses and responses to treatment^7,14,18,19^. Subtypes can sometimes be discovered from a single data type^7^, but often cannot be precisely defined without considering multiple layers of biological information^14^.

The classical approach to subtype discovery is to apply unsupervised learning methods, like hierarchical or centroid clustering, to a subset of input data that varies substantially between individuals. These input data can be binary (*e.g*. single nucleotide variants, SNVs), categorical (*e.g*. copy number alterations, CNAs), continuous (*e.g*. mRNA abundance), bounded continuous (*e.g*. methylation β-values, ranging from 0-1) or have other distributional features. Much molecular data is gene-based, but some represents processes like pathway activity or trinucleotide mutational signatures^20^. This classic approach has several limitations when applied to multiple data types simultaneously. First, standard unsupervised learning methods can produce artifactual results when applied to datasets with highly-variable distributional features, often implicitly assigning heavier weights to data types with many features or larger numerical ranges. To address this, some integrative subtyping algorithms transform input into a latent variable space while others use summary features from each individual data type^21–23^. Second, clinical practice routinely produces partial information, and most unsupervised learning methods struggle to accommodate large amounts of missing data^24,25^. Third, most existing methods do not exploit differential covariance or correlation across data types, nor provide clear understanding of how each data type contributes to the final subtyping.

We therefore created iSubGen (<u>i</u>ntegrative <u>sub</u>type <u>gen</u>eration) to create subtypes by directly quantifying inter-relationships between different data types. iSubGen recapitulates known molecular and histologic subtypes, robustly handles missing data, supports high subtype- and feature-number and seamlessly integrates gene-based and non-gene-based features.

## Results

### Development dataset

To develop iSubGen we first used the 1,991-patient METABRIC breast cancer dataset, which has clinical, CNA, SNV, miRNA abundance and mRNA abundance data, with the latter computationally deconvolved into tumour cell (TC) and tumour adjacent cell (TAC) components^14,26–28^. We initially focused on the 1,071 patients with complete data, and split these into the 684-patient training and 367-patient testing cohorts as in the original publication^14^. Initial method development used the 684 training cohort patients with complete data.

### Consensus integrative similarities

Typical approaches to subtype-identification quantify the relationship between each pair of patients using a similarity metric. For an *n*-patient cohort, this information is encoded in an *n* x *n* similarity matrix, which can be clustered using unsupervised machine-learning^29^. Thus clustering of CNA profiles (**Figure 1A**) generates CNA subtypes (**Figure 1B**) and clustering of SNV profiles (**Figure 1C**) generates SNV subtypes (**Figure 1D**) in the METABRIC training dataset.

**Figure 1.**
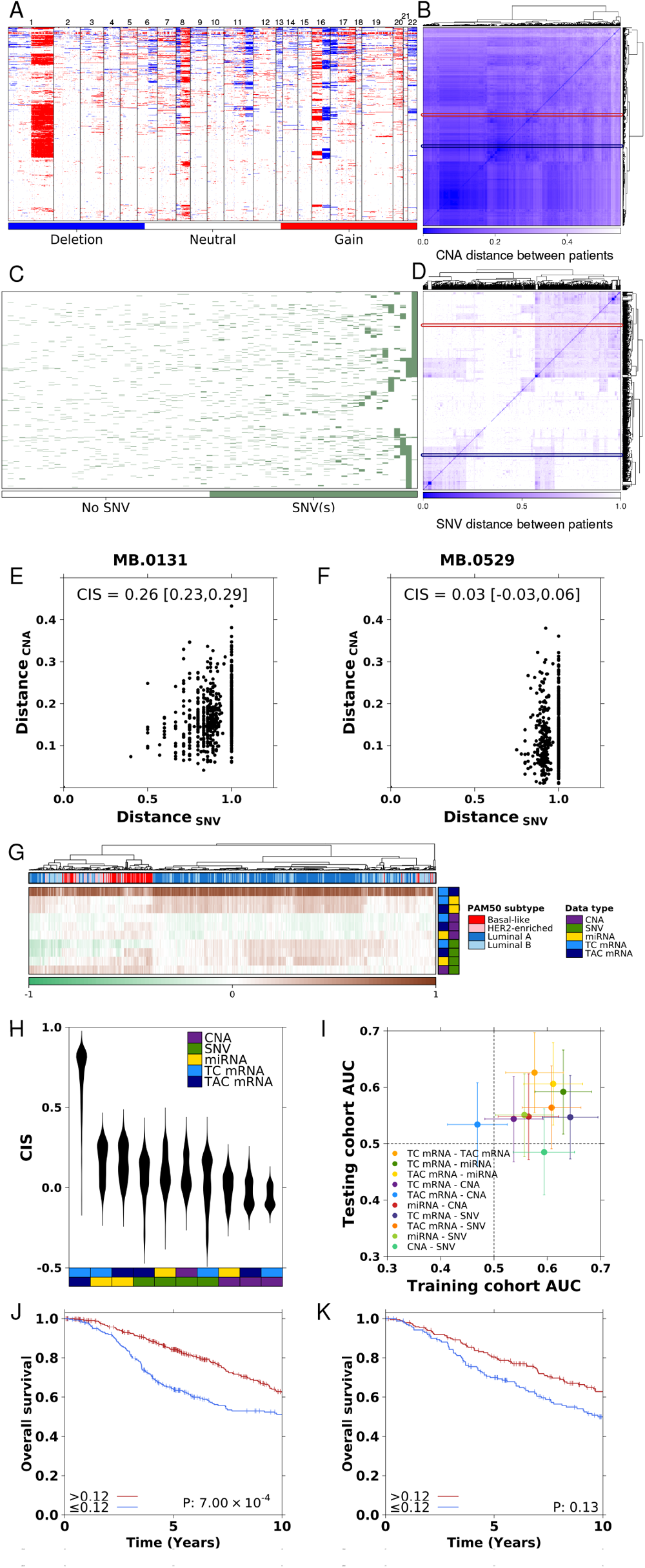
Integrative similarities. Pairwise integrative similarities in the training cohort of METABRIC breast cancer patients. (A) CNAs with genes ordered by genomic position on the x-axis and patients on the y-axis. Gains are red and deletions are blue. (B) CNA patient by patient similarity matrix using Jaccard distance as the similarity metric. SNVs for genes mutated in more than 13 patients. Genes (x-axis) are ordered by mutation frequency. Patients are on the y-axis. (D) SNV patient by patient similarity matrix calculated using SNVs without patient recurrence filtering and Jaccard distances. (E,F) Comparison of CNA and SNV similarities relative to patient MB.0131 (E) and MB.0529 (F). The CNA and SNV profiles for MB.0131 and MB.0529 are indicated on A and B by the boxes and arrows in red and blue respectively. Jaccard distances are used for measuring similarity in both CNA and SNV. MB.0131 was randomly selected as an example of a patient with positive CIS. MB.0529 was randomly selected as an example of a patient with a CIS near zero. (G) Patients grouped by clustering CISs. (H) The distributions of CIS for each data type pair. (I) Area under the receiver operator characteristic curve for predicting overall survival at five years using CISs. Error bars represent the 95% confidence intervals. (J,K) Overall survival differences for patients dichotomized by CIS_*T ACmRNA−miRNA*_ at the maximum geometric mean of the true positive rate and the false positive rate in the training cohort (I) and using the training cohort threshold in the testing cohort (J). P-values are from log-rank tests.

To integrate multiple data types into subtyping, there are two basic strategies. First, all data can be standardized to a common scale and a single metric applied to the appended matrix. Thus for an *n*-patient dataset with *m* data types each having *p*_*m*_ features, this results in performing similarity calculations on an *n* x Σ*p*_*m*_ feature matrix, producing a final *n* x *n* similarity matrix. This approach intrinsically preferences data types with more features or larger values because they hold more weight in similarity calculations^30^. An alternative strategy instead analyzes each data type separately and relates the *m* separate *n* x *n* similarity matrices. For example, each data type can be clustered separately, and then the patient classifications from each data type can themselves be clustered^11^. This discretizes patient classifications and intrinsically weights each data type either equivalently if cluster-number is held constant, or as a function of cluster-number if it is not.

To create a more flexible method of merging multiple data types, we directly reduced the pair of *n* x *n* similarity matrices for two data types into a continuous value representing the similarity between any two patients’ similarity profiles (**Figure 1E-F)**. Thus the two *n* length similarity vectors for a single patient, one per data type, are collapsed into a single value. Here, we used Spearman’s correlation to measure similarity. We used re-sampling to robustify this value, leading to a <u>c</u>onsensus integrative <u>s</u>imilarity (CIS) for each patient (**Figure 1E-F**). This yields a vector of *n* CISs for each pair of data types, yielding an *n* x [*m* x (*m* – 1) / 2] matrix encompassing the inter-relationships between data types for each patient. **Figure 1G** shows this matrix for 684 patients from METABRIC dataset, with simple unsupervised clustering applied to it. It is immediately apparent that luminal breast cancers cluster together and basal-like breast cancers cluster together, suggesting that CIS values reflect known disease subtypes.

CIS values are near zero for data types with independent (orthogonal) information, positive for data types with shared information and negative when patients similar to one another in one data type are dissimilar in the other. In METABRIC the median CIS across all data types was near zero (**Figure 1H**; median: 0.06, range: -0.38 to 0.88), with the most shared information between tumour cell and tumour adjacent cell (stromal) mRNA abundance (median CIS_TC-TAC_ = 0.77, range -0.07 to 0.88). The relationships between different types of information encapsulated in CISs were predictive of clinical features. In the training cohort, four of ten CISs predicted five-year survival (AUROC > 0.6) without applying any statistical learning, as validated in the 367-patient testing cohort (**Figure 1I)**. For example, stronger associations between TAC mRNA and miRNAs were associated with improved overall survival in training and testing cohorts (**Figure 1J-K**).

To further test the validity of CISs, we evaluated if CIS constituting mRNA abundance retained key information such as mRNA based subtypes of breast cancer (PAM50) and if other CISs were also predictive of breast cancer molecular subtypes. Using the training and testing cohorts, we compared CIS distributions between PAM50 subtypes (**Supplementary Figure 1A**). Almost all CISs differed amongst PAM50 subtypes (19/20, ANOVA q < 0.05). A random forest trained using CIS values predicted subtypes with AUROCs in the testing cohort ranging from 0.58 for luminal B to 0.95 for luminal A (**Supplementary Figure 1B**). CIS_TC mRNA-miRNA_ was the most important feature for the random forest luminal A classifier followed closely by CIS_TC mRNA-TAC mRNA_ and CIS_TAC mRNA–miRNA_ (**Supplementary Figure 1C**). We show CIS constituting mRNA abundance do indeed retain key information to predict mRNA based subtypes of breast cancer. We can extrapolate that other CISs may also be meaningful for novel subtypes.

To generalize this association of CISs with subtypes to other cancer types, we exploited TCGA data. We created pan-cancer training and testing cohorts each comprising 1,709 patients from twelve cancer types, and with six data types per patient (mRNA abundance, miRNA abundance, methylation, CNAs, SNVs and SNV trinucleotide signatures). All thirty CIS combinations in the training and testing cohorts distinguished cancer types (ANOVA q < 0.05; **Supplementary Figure 1D**). CIS distributions for some cancer types were bimodal, such as thyroid cancer (THCA) CIS_mRNA-SNV_. Histopathologic subtypes may cause this bimodality: in THCA, patients with tall cell thyroid cancer had higher CIS_mRNA-SNV_ than those with follicular thyroid cancer (Wilcoxon p = 3.09 × 10^−3^; **Supplementary Figure 1E**). Bimodality and high variance in CIS across many cancers increases the chance of finding subpopulations/subtypes. Random forest classifiers trained on CISs predicted all cancer types with AUROC_testing cohort_ > 0.9 (**Supplementary Figure 1F**). CISs vary in importance for predicting cancer types, with different CISs being important in distinguishing each cancer type (**Supplementary Figure 1G**). For example, CIS_methylation–mRNA_ was most important to identify liver cancers (LIHC), CIS_mRNA-miRNA_ for predicting kidney clear cell cancers (KIRC) and CIS_SNV-mRNA_ for kidney papillary cancers. Thus CISs can distinguish histological cancer types and subtypes.

### iSubGen framework and integrative subtyping

CISs capture the changing relationships between different types of data. To integrate them with information present in patterns of a single data type, we created a second set of engineered features. This second set of features was generated by training an autoencoder for each data type, and using its bottleneck layer as the set of independent reduced features (IRFs). iSubGen is thus a four-step subtype generation framework: consensus pairwise similarity construction (CIS generation), data type independent feature reduction (IRF generation), weighting of features and unsupervised machine-learning (**Figure 2**). The CIS values represent how different data types interrelate (**Supplementary Figure 2A**) while the IRF values identify general patterns within each data type (**Supplementary Figure 2B**). This strategy helps to balance groups of engineered features so that their relative weights are not primarily a function of the total feature number. In step three of the subtyping framework, the user sets the weightings of CIS *vs*. IRF and merges the two feature sets to create the combined engineered feature matrix. This provides a parameterizable decision for users, that can optimize based on internal features (*e.g*. cluster silhouette profiles) or external ones (e.g. separation of meta-data). Finally, applying pattern discovery to the combined sets of engineered features generates the final iSubGen subtypes. Here, we performed pattern discovery using consensus clustering^29^, but iSubGen supports multiple algorithms at each step. For example, CISs can use different correlation metrics or mutual information, with or without sub-sampling.

**Figure 2.**
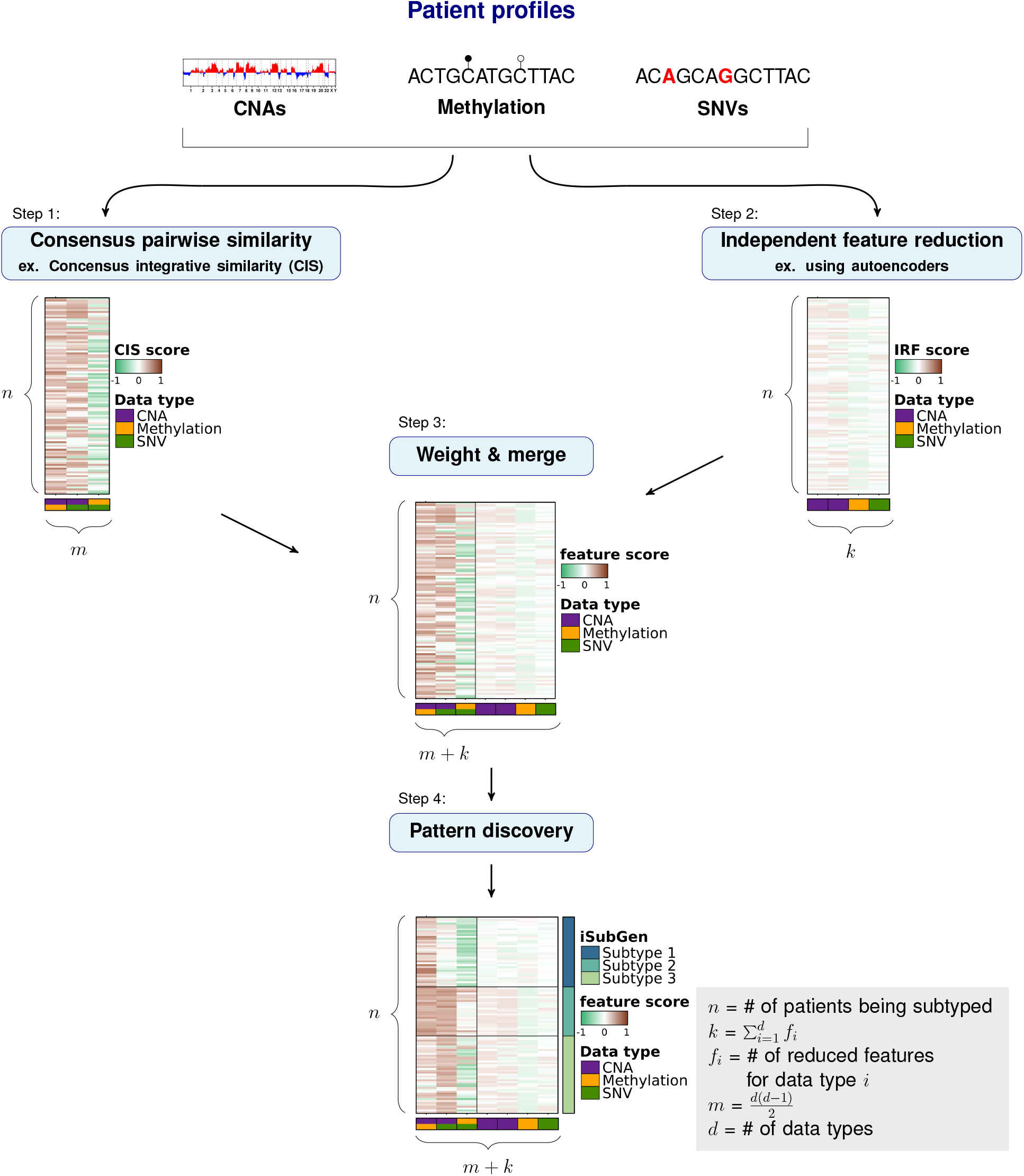
Integrative subtype generation overview. Schematic overview of iSubGen with three data types as an example for *n* patients. Each data type was separately run through feature reduction and combined in pairs for comparison of the patient profiles using similarity measures. Output from feature reduction (*n* rows by *k* columns) and similarity comparison (*n* rows by *m* columns) were rescaled and reweighted if necessary and merged into a single matrix for unsupervised machine learning to create the final classifications. We used the autoencoder bottleneck layer for independent feature reduction and CISs as our pairwise similarity measures.

### Pan-cancer grouping discovery with iSubGen

To demonstrate how iSubGen combines CISs and IRFs to generate robust subtypes, we applied it to the pan-cancer cohort evaluated above (**Supplementary Figure 1D-G**). Using six data types (miRNA abundance, mRNA abundance, methylation, CNA, SNV and trinucleotide signatures), we subtyped the two 1,709-patient subsets of twelve cancer types separately using iSubGen (**Supplementary Figure 3, Supplementary Table 7)**. Each subset was independently analysed to evaluate iSubGen subtype consistency. Comparing the adjusted Rand index of the iSubGen clusters with TCGA cancer types, we identified fourteen iSubGen groupings in both subsets: iSubGen-P1 through iSubGen-P14 and iSubGen-Q1 through iSubGen-Q14 in the discovery and validation cohorts respectively.

iSubGen-P1 which is comprised almost entirely of skin cutaneous melanoma (SKCM) had the highest CIS_SNV–signature_ (**Supplementary Figure 3A**). Lung adenocarcinomas (LUAD), stomach and esophageal carcinoma (STES) breast cancers (BRCA), bladder cancers (BLCA) and head and neck squamous cell cancers (HNSC) were classified together in multiple groups. Thyroid cancers (THCA) were separated from the other cancers into two thyroid cancer groups: iSubGen-P10/iSubGen-Q10 and iSubGen-P11/iSubGenQ11 (**Supplementary Figure 3B**,**D**). iSubGen-P10/iSubGen-P11 contained 90% (135/150) and iSubGen-Q10/iSubGen-Q11 contained 97% (146/150) of THCA patients in their respective cohorts. The obvious differences between these two thyroid cancer groups was that CIS_SNV– methylation_ (Wilcox p < 2.2 × 10^−16^), CIS_SNV-mRNA_ (Wilcox p < 2.2 × 10^−16^) and CIS_SNV-miRNA_ (Wilcox p < 2.2 × 10^−16^) were higher in iSubGen-P10/iSubGen-Q10 than iSubGen-P11/iSubGen-Q11, representing subtypes of thyroid cancer (**Supplementary Figure 3C**,**E**). The CIS values for iSubGen-P and iSubGen-Q groupings had high concordance (**Supplementary Figure 3F**). Thus iSubGen generates CIS and IRF values that are both useful for supervised learning and that allow unsupervised learning to independently create concordant classifications in two pan-cancer datasets.

### Integrative molecular- and pathway-based breast cancer subtyping

To demonstrate the utility of iSubGen for integrative multi-modal subtype discovery, we next applied it to the METABRIC breast cancer cohort, integrating 19,877 mRNA features for both TC and TAC mRNA, 18,852 CNA features, 823 miRNA features and SNV mutation status of 173 driver genes. When applied to these five data types, iSubGen identified five subtypes (**Supplementary Figure 4A, Supplementary Table 1**), which differed in patient survival (**Supplementary Figure 4B**). We named subtypes such that patients in iSubGen-B1 having the best outcome and iSubGen-B5 the worst. iSubGen-B1 and iSubGen-B2 had lower tumour grade (**Supplementary Figure 4C**) and size (**Supplementary Figure 4D**) than iSubGen-B4 and iSubGen-B5. The five iSubGen-B subtypes were tightly associated with the PAM50 subtypes^7,31^. Notably iSubGen-B5 contained most HER2-enriched and basal-like breast cancers in both training and testing cohorts (**Supplementary Figure 4E**), linked to its lower CIS_TC mRNA – TAC mRNA_ (Wilcox p < 2.2 × 10^−16^), CIS_TC mRNA - miRNA_ (Wilcox p < 2.2 × 10^−16^) and CIS_TAC mRNA – miRNA_ (Wilcox p < 2.2 × 10^−16^) (**Supplementary Figure 4F**). This reflects a higher transcriptome similarity amongst luminal breast cancers than amongst HER2-enriched or basal-like ones. Even amongst the luminal cancers, the good outcome iSubGen-B1 and iSubGen-B2 subtypes had higher CIS_SNV-mRNA_ and CIS_SNV-miRNA_ (p < 2.2 × 10^−16^; **Supplementary Figure 4F**). We similarly identified strong associations between iSubGen-B and METABRIC IntClust subtypes^14^ (**Supplementary Figure 4G**). Overall, the iSubGen-B subtypes showed concordance with the known subtypes, and identified some novel groupings and highlighted differences in the covariance structure of different types of molecular data across subtypes.

To demonstrate that iSubGen can be useful with only a single molecular data type, we next focused on the mRNA abundance data of METABRIC, evaluated as a set of 13 cancer hallmark pathways^32,33^ (**Supplementary Figure 5A, Supplementary Table 2**). In the training cohort iSubGen identified seven hallmark subtypes, associated with overall survival (**Supplementary Figure 5B**). The CISs between cancer hallmarks were generally higher (median CIS 0.5) than CISs between different data types (median CIS 0.06, Wilcox p < 2.2 × 10^−16^). iSubGen hallmark-based breast cancer subtypes (iSubGen-H1 through iSubGen-H7) were associated with PAM50 (**Supplementary Figure 5C**), IntClust (**Supplementary Figure 5D**) and iSubGen-B subtypes (**Supplementary Figure 5E**). In general, higher CISs between hallmarks in the iSubGen-H subtypes associated with better overall patient survival (**Supplementary Figure 5A-B**). iSubGen subtypes are concordant with the idea that tumours with more dysregulation across data types and signaling pathways have poorer outcome.

The iSubGen-B and iSubGen-H subtypes are assessing breast cancer by two different paradigms: iSubGen-B is a genome-wide approach and iSubGen-H is a pathway approach. We combined the engineered features from these two approaches within a single model to demonstrate the flexibility of iSubGen. Together, these six molecular data types comprise 39,725 molecular features and 13 pathway activities. We identified ten subtypes in our training cohort (**Figure 3A, Supplementary Table 3**) and again named them by their association with overall survival: iSubGen-BH1 through iSubGen-BH10 (**Figure 3B**). The iSubGen-BH subtypes associated with PAM50 subtypes and improved separation of basal-like breast cancers and HER2-enriched cancers relative to iSubGen-B and iSubGen-H (**Figure 3C**). These associations validated in the testing cohort *via* centroid classification (**Figure 3D**), highlighting the reproducibility of iSubGen subtypes.

**Figure 3.**
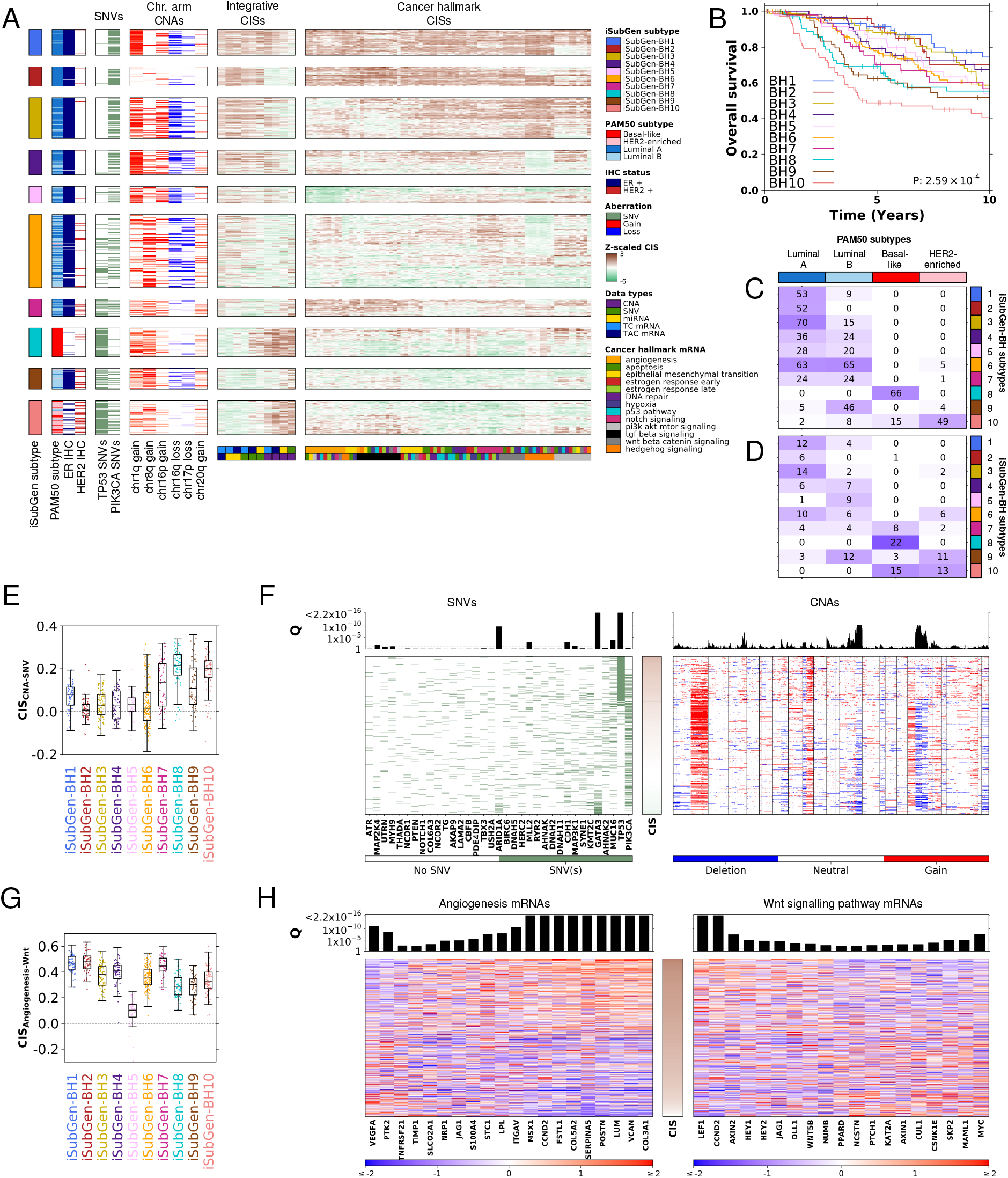
Breast cancer iSubGen combining integrative omics features and cancer hallmark mRNA features. (A) Using the iSubGen, the breast cancer patients in the training cohort were classified into ten subtypes using integrative -omics features and mRNA cancer hallmark features. (B) Overall survival for the iSubGen-BH breast cancer sub-types. P-value is from a log-rank test. (C) Comparison of the iSubGen breast cancer subtypes and PAM50 subtypes in the training cohort. Heatmap colouring represents the number of the patients in each overlap. (D) Comparison of the iSubGen breast cancer subtypes and PAM50 subtypes in the testing cohort. (E) Comparison of CIS_*CNA−SNV*_ distributions between iSubGen-BH subtypes in the training cohort. (F) Association of SNV and CNA features with the CIS_*CNA−SNV*_ . The top barplot shows significance between each feature and CIS_*CNA−SNV*_ using Wilcoxon rank sum tests and FDR-adjustment. (G) Comparison of CIS between angiogenesis mRNA set and Wnt/-catenin signaling mRNA set for iSubGen-BH subtypes in the training cohort. (H) Association of CIS_*angiogenesisW nt/−cateninsignaling*_ with z-scaled mRNA abundance from each gene set (q *<* 0.01). The top barplot shows significance between each mRNA and the CIS using Spearman’s correlation and FDR-adjustment. Genes are ordered by Spearman’s correlation with correlations decreasing out from middle and the CIS panel.

To characterize the iSubGen-BH CISs, we examined their associations with the individual input features. Higher CIS_CNA–SNV_ was associated with iSubGen-BH7 (p = 1.4 × 10^−7^), iSubGen-BH8 (p < 2.2 × 10^−16^), iSubGen-BH9 (p = 1.7 × 10^−7^) and iSubGen-BH10 (p < 2.2 × 10^−16^) compared to iSubGen-BH1 through iSubGen-BH6 (**Figure 3E**). We identified six SNV associations (of 156) and 1,283 CNA associations (of 10,662; q < 0.01) where mutation of a specific gene associated with higher or lower CIS_CNA-SNV_ (**Figure 3F**). Patients with *TP53* SNVs had high CIS_CNA-SNV_, while *GATA3* SNVs and *ARID1A* SNVs were associated with low CIS_CNA-SNV_. Amongst the associated CNAs, deletion of the q arms of chromosomes 11 and chromosome 16 were associated with lower CIS_CNA-SNV_. We also examined individual input features association with the hallmark CISs. Lower CIS_angiogenesis – Wnt/β-catenin_ differentiated iSubGen-BH5 from the other iSubGen-BH subtypes (Wilcox p < 2.2 × 10^−16^, **Figure 3G**). There was 36 angiogenesis mRNAs and 42 Wnt/β-catenin signaling mRNAs in the individual hallmark gene sets that were used to calculate CIS_angiogenesis – Wnt/β-catenin_. We found three genes, including *VEGFA*, from the angiogenesis gene set and ten genes, including *MYC*, from the Wnt/β-catenin signaling gene set where higher mRNA abundance was associated with lower CIS_angiogenesis – Wnt/β-catenin signaling_ (Spearman q < 0.01; **Figure 3H**). There were seventeen genes from the angiogenesis gene set and nine genes from the Wnt/β-catenin signaling gene set where lower mRNA abundance was associated with lower CIS_angiogenesis – Wnt/β-catenin signaling_ (Spearman q < 0.01). Thus iSubGen enhances subtype development by integrating individual features (IRFs) with feature-feature interactions (CISs).

### Subtyping using genic and non-genic molecular data

To evaluate whether iSubGen could be used to generate subtypes from more diverse molecular data, we sought to subtype using a combination of gene-based and mutational process information. We used trinucleotide signatures^20^ for 557 patients from three kidney cancer types from the pan-cancer TCGA datasets^34–36^, which to our knowledge have not been previously integrated into multi-modal subtyping strategies. Each cancer type had six available data types: CNA, SNV, trinucleotide signature exposures, methylation, mRNA abundance and miRNA abundance. We randomly divided patients with all six data types into equal-sized training and testing cohorts. In the training cohort, we used iSubGen to identify eight subtypes: iSubGen-K1 through iSubGen-K8 (**Figure 4A, Supplementary Table 4**). iSubGen-K1 contained almost all kidney chromophobe (KICH) cancers, while iSubGen-K2 through iSubGen-K5 comprised predominantly kidney papillary (KIRP) cancers and iSubGen-K6 through iSubGen-K8 predominantly clear cell (KIRC) cancers. Centroid classification in the testing cohort validated the presence and relative frequencies of these subtypes (**Figure 4B**). Interestingly, KIRC patients in iSubGen-K5 had poorer overall survival than other KIRC patients in both training and testing cohorts (**Figure 4C**), while KIRP patient survival was not associated with iSubGen-K subtypes (**Figure 4D**). Trinucleotide signatures were associated with both histological classifications^20^ and iSubGen-K subtypes (**Figure 4E**). KIRC patients classified in iSubGen-K6 and iSubGen-K8 were associated with a lower exposure of patients to SBS5, which has unknown etiology. iSubGen-K4 and KIRC classified patients were associated with a higher exposure to SBS5. SBS13 and SBS2, which are attributed to AID/APOBEC activity, were associated with KIRP patients. Thus iSubGen provides a framework for integrating both mutational and mutational-process information into subtype discovery.

**Figure 4.**
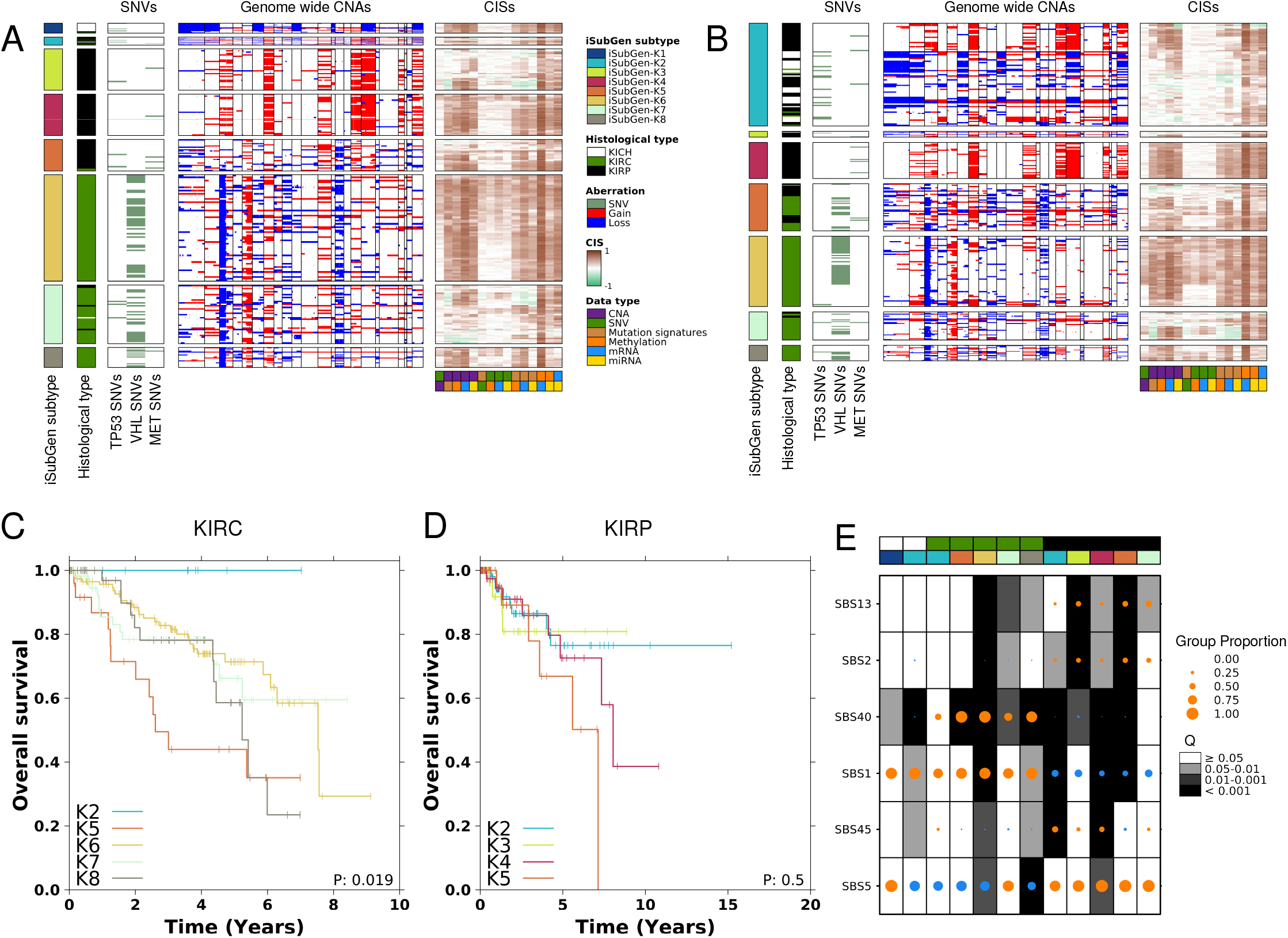
Kidney cancer iSubGen using non-gene-based features. (A) Using iSubGen, patients in the training cohort were classified into six subtypes. (B) Centroid classification of the iSubGen-K subtypes in the test cohort. (C) Overall survival between iSubGen classifications for KIRC patients. Groups with less than 10 patients are not included. (D) Overall survival between iSubGen classifications for KIRP patients. Groups with less than 10 patients are not included. P-values are from log-rank tests. (E) Association of patients in the training and test cohorts with trinucleotide mutation signatures. Each column is a group of patients. The top covariate shows the TCGA histological cancer type and the second covariate is the iSubGen classification of the patients. Patient groups with less than ten patients are not shown. Each dot is sized to proportion of patients with mutations from the trinucleotide signature. If the dot is orange, then the proportion for the group is greater than the proportion in the patients not in the group. Similarly if the dot is blue, then the proportion is less than in the other patients. The background shading is the q-value from the proportion test comparing the proportion for patients in the group to the proportion for those not in the group.

### iSubGen subtyping is robust to missing data

Because human cancers vary in size, and are often profiled from biopsy rather than surgical specimens, it is common for only a subset of possible molecular assays to be performed. For this, and many other reasons, missing data is common in genomic studies. To evaluate iSubGen’s performance in the face of missing data, we randomly separated TCGA lung cancer data into a 512 patient training cohort and a 509 patient testing cohort^37,38^. Overall 446/1021 patients (47%) lacked one or more of the six data types used in classification, split evenly between training and testing cohorts (**Figure 5A, Supplementary Table 5**). To select the number of iSubGen subtypes, we assessed the association between histological subtypes and different numbers of iSubGen clusters using the adjusted Rand index in the training cohort. Lung cancers formed two subtypes: iSubGen-L1 and iSubGen-L2. Subtype structure was robust to missing data (**Figure 5B, Supplementary Table 5**). iSubGen-L1 largely comprised lung adenocarcinomas and iSubGen-L2 largely comprised lung squamous cell carcinomas in both training (**Figure 5C**) and testing cohorts (**Figure 5D**). Overall 89% (230/257) of training and 87% (227/260) of testing cohort lung adenocarcinomas were in iSubGen-L1. Similarly 87% (221/255) of training and 80% (198/249) of testing cohort lung squamous cell carcinomas were in iSubGen-L2. iSubGen-L2 had higher median CIS_mRNA-SNV_ than iSubGen-L1 (median_training L1_=-0.04, median_training L2_=0.18 P_training_ < 2.2 × 10^−16^; median_testing L1_=-0.04, median_testing L2_=0.16, P_testing_ < 2.2 × 10^−16^; **Figure 5E-F**). To visualize these underlying associations between data types, we focused on CISs from two exemplar patients and on mRNA and SNV features prior to feature engineering. TCGA.44.5645 had lower overall CIS_mRNA-SNV_ and near-median values for iSubGen-L1 patients (**Figure 5G**). TCGA.78.7155 had the highest CIS_mRNA-SNV_ of all patients in the training cohort and had high CIS_mRNA-SNV_ for iSubGen-L2 (**Figure 5H**). TCGA.44.5645 and TCGA.78.7155 both cluster with their respective histological subtype using mRNA abundance (**Figure 5I**). By contrast, SNVs did not separate patients by histological subtype (**Figure 5J**): TCGA.44.5645 has more total somatic SNVs than TCGA.78.7155 and clusters with other highly mutated tumours. The low CIS_mRNA-SNV_ values show that SNVs and mRNA provide orthogonal information – leading iSubGen to create composite subtypes that merge them.

**Figure 5.**
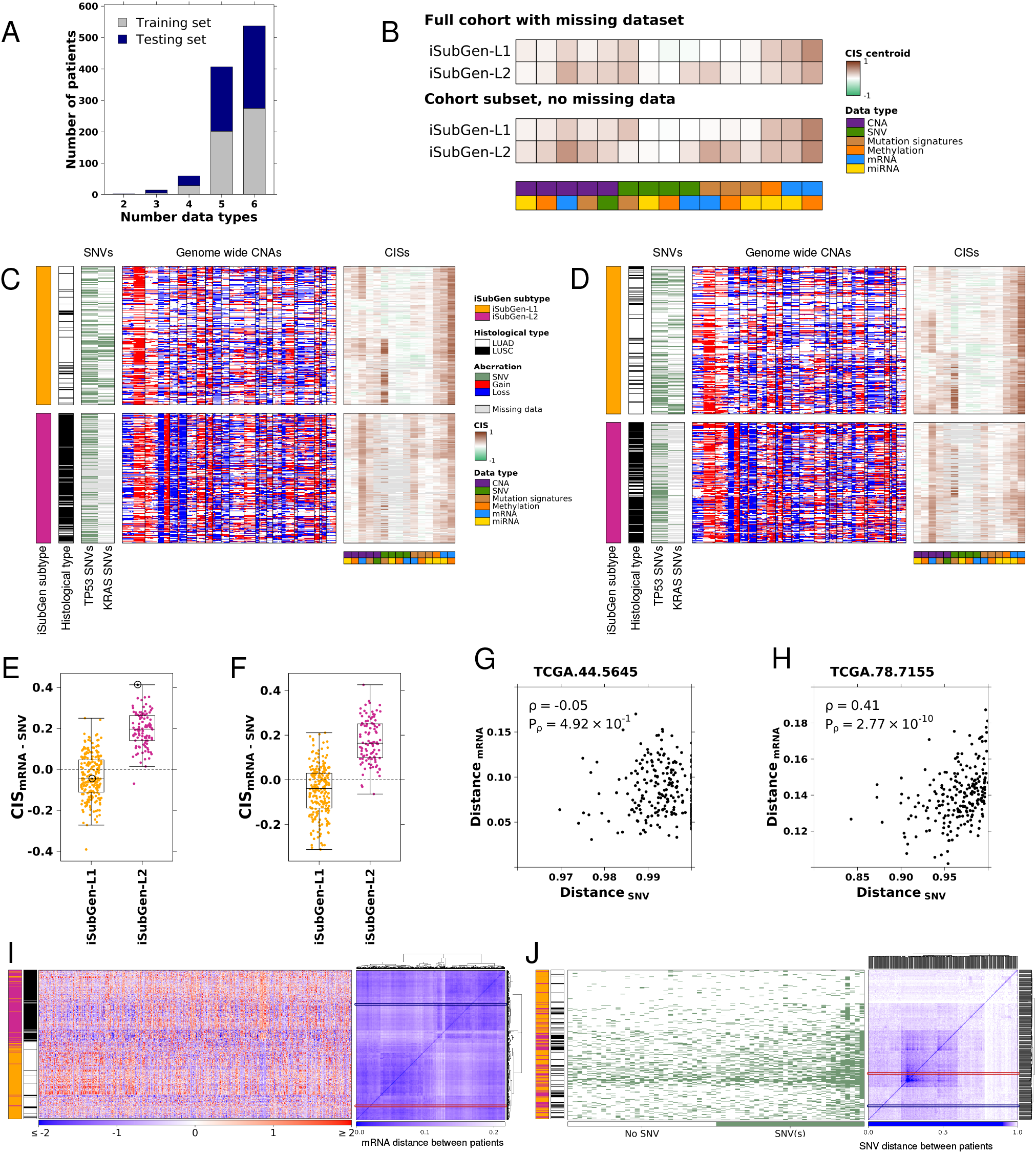
iSubGen is robust to missing data in lung cancer. (A) The number of data types that each patient has. (B) Using iSubGen, patients from the training cohort were classified into two subtypes including patients with missing data. The top panel is the centroids from subtyping with missing data. To assess the effect of missing data, a subset of patients that had all six data types were also independently clustered and these centroids are in the bottom panel. (C) iSubGen classification of the training cohort. (D) Centroid classification of the iSubGen-L subtypes in the testing cohort. (E) CIS_*mRNA−SNV*_ for training cohort with TCGA.78.7155s and TCGA.44.5645s CISs circled. (F) CIS_*mRNA−SNV*_ for testing cohort. (G) The CIS for patient TCGA.44.5645 which is the patient with the median CIS_*mRNA−SNV*_ in iSubGen-L1. (H) The CIS for patient TCGA.78.7155 which is the patient with the highest CIS_*mRNA−SNV*_ in iSubGen-L2. (I) Z-scaled mRNA abundance for mRNAs with standard deviation greater than 2. mRNA on the x-axis and patients on the y-axis. Far right plot, mRNA abundance patient by patient similarity matrix using 1 - Pearson correlation as the similarity metric. (J) SNVs for genes mutated in more than 50 patients. Genes (x-axis) are ordered by mutation frequency. Patients are on the y-axis. Far right plot, SNV patient by patient similarity matrix calculated using SNVs without patient recurrence filtering and Jaccard distance as the similarity metric. TCGA.78.7155 is circled in blue and TCGA.44.5645 is circled in red on I and J.

To assess the association of CIS values with epidemiologic features we considered sex differences, which have been widely reported in lung cancer^39–41^. We tested whether CISs differed between tumours arising in patients with XX and XY germline chromosome conformations; 7/15 CISs were associated with sex (**Supplementary Table 6**). For example, XY patients had higher CIS_mRNA-SNV_ than XX patients: mRNA and SNV profiles were more concordant in lung tumours arising in men than those arising in women. Thus CISs reflect underlying epidemiologic features, independent of missing data.

## Discussion

Many factors influence the status and progression of cancer: somatic mutations (CNAs, SNVs, *etc*.), epigenomic alterations, chromatin reorganization and external cellular signals all occur on a background of the individuals’ health, stress and exposures over a lifetime. Capturing how all these data types interrelate is an open problem, subject to intensive investigation. We introduce iSubGen to capture the associations between different types of data. iSubGen uses a novel metric called CIS to capture how different data types interrelate. If feature patterns from two data types define the same patient associations (high CIS), then the two data types may reflect a regulatory relationship of some type. For example, a higher CIS_SNV-mRNA_ could be explained by a set of mutated genes, such as transcription factors, that drive broad mRNA abundance patterns. CISs, along with a reduced form of the single data type information termed IRFs, both serve as intermediaries for integrative subtype discovery and can be used for direct supervised biomarker development.

iSubGen facilitates maximum data type inclusion and modular replacement of the framework steps to personalize for different situations. However, with increased options comes increased parameterization and the need to check that the underlying engineered features are reproducible. Indeed iSubGen does not directly incorporate prior information (although its robustness to missing data provides a natural pathway for doing so). Almost all subtype-development approaches face this challenge: there is no one metric to quantitatively optimize clustering results when selecting weightings, the number of clusters and the ultimate subtypes. We considered inter-subtype differences, association with CISs, prognostic associations and, since we were assessing well characterized cancer types, association with known histopathologic subtypes. In any subtyping, it is up to the user to decide what is most important in choosing a subtype number when multiple different values can bring statistically reasonable results using domain knowledge.

A caveat of iSubGen is its relative execution time to other subtype discovery methods. The calculation of CIS values is computationally demanding because of the number of similarity calculations for, first, similarity matrices for each data type and second, CIS, calculations between data types. CIS is a consensus metric so requires iterations of all of the similarity calculations. For large numbers of patients, like our pan-cancer analysis, creation of the CIS matrix can take several hours on a single CPU (one core), although this can be readily parallelized.

Subtypes provide fundamental understanding about polygenic disease – they identify groups of patients whose current disease appears similar, and thus might share both similar histories and future responses to treatment. Potential applications of iSubGen extend to almost any complex biological system. For example, to understand the effect of human microbiomes on health, we will need to recognize patterns across underlying human genetics, epigenetics, metabolomics and the microorganisms present in the gut. Data arising from mobile devices provide a completely different setting with a plethora of data types to combine and interpret. iSubGen provides a flexible framework to capture multi-modal interactions in diverse data science applications.

## Methods

### METABRIC breast cancer dataset

The METABRIC cohort contains 1,991 patients each with a primary fresh frozen breast cancer specimen^14,26,27^. METABRIC annotation includes overall survival and PAM50 subtype classifications. Six patients had subtype classification NC (not classified). The METABRIC dataset includes mRNA abundance, CNAs, miRNA and SNVs. The relative mRNA abundances of 19,877 genes were profiled using Illumina HT-12 v3 microarrays for 1,988 patients. CNA data covers 18,852 genes profiled using Affymetrix SNP 6.0 microarrays for 1,989 patients. There are 823 relative miRNA abundances profiles using Agilent Human miRNA Microarray 2.0 for 1,285 patients^26^. The METABRIC cohort also included targeted sequencing data covering 173 genes frequently mutated in breast cancer (*i.e*. candidate driver genes) with somatic SNV calls^27^. The mRNA abundance was deconvolved into tumour cell and tumour adjacent cell mRNA abundance using ISOpure^28,42,43^. TC/TAC deconvolution was performed for all patients in the training cohort and all patients in the testing cohort. We used the training and testing cohort divisions from the METABRIC paper. Subtypes were discovered using only the training cohort.

### TCGA data

TCGA datasets were downloaded from Broad GDAC Firehose (https://gdac.broadinstitute.org/), release 2016-01-28. We used the mRNA abundance, CNAs, SNVs for the TCGA samples. The mRNA abundances of 20,531 genes were profiled using exome sequencing. CNA data covers 24,776 genes profiled using Affymetrix SNP 6.0 microarrays. There are 18,152 genes with a mutation for the SNV data. Per patient trinucleotide mutational signatures calls were also downloaded. miRNA was downloaded from the GDC Data Portal (https://portal.gdc.cancer.gov/), data Release 25.0 – July 22, 2020. There were 1,881 miRNA abundance profiled through sequencing.

For the kidney cancer cohort, we used 241 kidney renal papillary cell carcinomas (KIRP), 267 kidney renal clear cell carcinomas (KIRC) and 49 chromophobe renal cell carcinomas (KICH)^34–36^. We only used patients with all six data types. Patients were randomly divided per subtype to create equally sized training and testing cohorts.

For the lung cancer cohort, we used 1,021 patients from the TCGA lung cancer cohorts^37,38^, which is a combination of lung adenocarcinoma (LUAD) and lung squamous cell carcinoma (LUSC). Using random sampling per subtype, we divided the patients into a training cohort of 512 and a testing cohort of 509 patients.

For the pan-cancer cohort, twelve TCGA datasets with more than 200 patients with the data types were chosen: BLCA, BRCA, HNSC, KIRC, KIRP, LGG, LIHC, LUAD, PRAD, SKCM, STES, THCA. From each cancer type, we equally divided up to 300 randomly selected patients in two non-overlapping subsets. In total each subset had 1709 patients.

### Survival associations of CISs

We created a receiver operator curve using the CISs for each pair of data types. For each pair of data types, the CISs were dichotomized at every possible threshold and agreement with overall survival was assessed. For further examples of the survival associations, we created Kaplan-Meier curves and assessed the survival association using log-rank tests for CIS_TAC mRNA – miRNA_. CIS dichotomization threshold was chosen to maximize the harmonic mean of true positive and false positive rates for predicting five year overall survival using all the patients in the training cohort.

### Random Forest classifiers

We created random forest classifiers predicting breast cancer subtype or pan-cancer cancer type from CISs using the randomForest (v4.6-14) R package.

### Independent reduced features

The reduced feature matrix is a matrix where each row is a patient and each column is an IRF. For each data type, an autoencoder was created using the keras (v2.1.5) and tensorflow (v1.10) packages in R. RNA profiles were scaled before inputting to the autoencoder. The autoencoders were trained with mean squared error loss function, Adam optimization and tanh as the activation function. Each autoencoder had three hidden layers with fifteen nodes, two nodes, fifteen nodes respectively. We tested one, three and five hidden layers with various node sizes (1, 2, 5, 15, 30, 25, 50, 100, 200). The IRFs were then extracted from the layer with the bottleneck layer (here the layer with two nodes). These IRFs for each data type were combined into a matrix where each column corresponded to a node in the bottleneck layer from the autoencoders. There were two columns from each data type.

### Consensus integrative similarities

The pairwise comparison matrix is a matrix where each row is a patient and each column is a pair of data types. The entries in the matrix are correlations or consensus correlations. To calculate the correlation for a patient and a pair of data types, the similarities between that patient and each of the other patients in the cohort were calculated for each of the data types. These similarities were then correlated between two data types. This was repeated for each patient and each pair of data types using Spearman’s correlations. The similarity metric varied depending on the molecular data type. For CNA, SNV, trinucleotide mutational signatures data types, we used Jaccard distance as the similarity metric. For mRNA, miRNA and methylation data types, we used 1 - Pearson’s correlation as the similarity metric. To create CISs, patients were correlated with bootstrapping and each CIS was the median correlation from the sub-sampled repetitions. For each bootstrap, 80% of the patients were sampled without replacement and all the patients were individually correlated to that 0.8 subset of patients. This was repeated 10 times and the median of the correlations for each patient and data type pair.

### Integrative Subtype Generation (iSubGen)

There are four steps to creating subtypes with iSubGen. (1) Create a pairwise comparison matrix which assesses the relationships between patient similarities in a pairwise approach such as CISs. (2) Create a reduced feature matrix to assess the main pattern of each independent matrix. (3) Combine the pairwise similarity matrix and the reduced feature matrix with appropriate re-weighting. (4) Perform pattern discovery on the combined matrix. We used consensus clustering (v1.8.1)^29^ with a seed of 17, with 1000 clustering repetitions and Euclidean distance metric and hierarchical clustering with Ward linkage. The number of clusters was determined using the consensus cluster results, including the consensus matrix and cumulative distribution functions and association with CISs and clinical features.

### Breast cancer integrative multi-omics subtypes

iSubGen was run on the 684 patients in the training cohort with all five data types: CNA, SNV, miRNA abundance, TC mRNA abundance and TAC mRNA abundance. TC mRNA, TAC mRNA and miRNA features were z-scaled per feature before autoencoder training. For each data type, the autoencoder had three hidden layers with fifteen nodes, two nodes, fifteen nodes respectively. Therefore there was two independent reduced features (IRFs) per data type. Consensus clustering was performed for 2 to 10 subtypes with 0.8 sub-sampling of features and patients. A weighting of 1:8 for CISs to independent reduced features was selected. Five subtypes was selected by visual assessment of iSubGen subtypes with CIS, PAM50 subtypes and prognosis. There were 367 patients in the testing cohort with all five data types. Testing cohort independent reduced features were created using the trained autoencoders with TC mRNA, TAC mRNA and miRNA features scaled using the mean and standard deviations from the training cohort. Testing cohort CISs were calculated for each patient relative to the patients in the training cohort, not relative to the patients in the other patients in the testing cohort.

### Breast cancer subtypes from mRNA of cancer hallmarks and pathways

iSubGen was run on the 996 patients in the training cohort with mRNA abundance. Thirteen gene sets from the MSigDB hallmark gene sets collection were selected and mRNA for the genes from each of these sets was used as a separate data type. mRNA features were z-scaled per feature per mRNA set before autoencoder training. For each data type, the autoencoder had three hidden layers with fifteen nodes, two nodes, fifteen nodes respectively. Therefore there was two IRFs per data type. Consensus clustering was performed for 2 to 10 subtypes with 0.8 sub-sampling of features and patients. A weighting of 1:4 for CISs to independent reduced features was selected. Nine subtypes was selected by visual assessment of iSubGen subtypes with CIS, PAM50 subtypes and prognosis.

### Breast cancer subtypes combining integrative -omics features and mRNA sets

iSubGen was run on the 684 patients in the training cohort with all five data types: CNA, SNV, miRNA abundance, TC mRNA abundance and TAC mRNA abundance. Features were used as described from breast cancer integrative multi-omics subtypes and breast cancer subtypes from mRNA of cancer hallmarks and pathways. A weighting of 1:2 for CISs to IRFs was selected. Consensus clustering was performed for 2 to 18 subtypes with 0.2 sub-sampling of features and 0.8 sub-sampling of patients. Ten subtypes was selected by visual assessment of iSubGen subtypes with CIS, PAM50 subtypes and prognosis.

### Kidney cancer subtypes

iSubGen was run on the 283 patients in the training cohort with all six data types: CNA, SNV, trinucleotide mutational signatures, methylation, mRNA abundance and miRNA abundance. mRNA and miRNA features were z-scaled per feature before autoencoder training. For each data type, the autoencoder had three hidden layers with fifteen nodes, two nodes, fifteen nodes respectively. Therefore there was two IRFs per data type. Consensus clustering was performed for 2 to 10 subtypes with 0.8 sub-sampling of features and patients. A weighting of 1:2 for CISs to independent reduced features was selected. Five subtypes was selected by visual assessment of iSubGen subtypes with CIS, TCGA kidney cancer types and prognosis. There were 274 patients in the testing cohort with all six data types that were classified using centroid classification. Testing cohort independent reduced features were created using the trained autoencoders with mRNA and miRNA features scaled using the mean and standard deviations from the training cohort. Testing cohort CISs were calculated for each patient relative to the patients in the training cohort, not relative to the patients in the other patients in the testing cohort.

### Lung cancer subtypes

iSubGen was run on the 512 patients in the training cohort with any of the six data types: CNA, SNV, trinucleotide mutational signatures, methylation, mRNA abundance and miRNA abundance. All patients had at least two data types. If missing data, NA was used in the matrix. mRNA and miRNA features were z-scaled per feature and trinucleotide mutational signatures features were log_10_-scaled before autoencoder training. For each data type, the autoencoder had three hidden layers with fifteen nodes, two nodes, fifteen nodes respectively. Therefore there was two IRFs per data type. Consensus clustering was performed for 2 to 10 subtypes with 0.5 sub-sampling of features and patients. Diana, instead of hclust, was used within consensus clustering because it can handle clustering missing data. A weighting of 1:8 for CISs to independent reduced features was selected. Two subtypes were selected looking at association with TCGA lung cancer type while also minimizing the number of clusters.

Using a subset of 126 patients (63 LUAD, 63 LUSC) with all six data types, we ran iSubGen as we did with the cohort including patients with missing data. Since the cohort with missing data has equivalent numbers of LUAD and LUSC patients, we downsampled to have a cohort with equal proportion of each subtype with all data types. There were 63 LUSC patients with all data types so we randomly selected 63 LUAD patients from the 212 LUAD patients with all data types. There were 279 patients in the testing cohort with all five data types that were classified using centroid classification. Testing cohort independent reduced features were created using the trained autoencoders with mRNA and miRNA features scaled using the mean and standard deviations from the training cohort and trinucleotide mutational signatures features were again log_10_-scaled. Testing cohort CISs were calculated for each patient relative to the patients in the training cohort, not relative to the patients in the other patients in the testing cohort.

### Pan-cancer subgroups

Two subset were randomly created with the twelve TCGA datasets (BLCA, BRCA, HNSC, KIRC, KIRP, LGG, LIHC, LUAD, PRAD, SKCM, STES, THCA) with more than 200 patients for the six data types (mRNA, CNA, SNV, trinucleotide mutational signatures, methylation, miRNA). From each cancer type, we randomly selected 300 patients or all the patients and evenly divided them between the two subsets. There was 1,709 patients in the first subset and 1,709 patients in the second subset. Both subsets were independently run through iSubGen. mRNA features were z-scaled per feature before autoencoder training. For each data type, the autoencoder had three hidden layers with fifteen nodes, two nodes, fifteen nodes respectively. Therefore there was two IRFs per data type. Consensus clustering was performed for 2 to 30 subtypes with 0.8 sub-sampling of features and 0.1 sub-sampling of patients. A weighting of 1:4 for CISs to independent reduced features was selected. Fourteen subtypes was selected in both subsets by assessing association of the iSubGen groups with cancer types using adjusted Rand index.

### Visualization

All plotting was performed in the R statistical environment (v3.4.3) using the lattice (v0.20-38), latticeExtra (v0.6-28), RColorBrewer (v1.1-2) and cluster (v2.0.7-1) packages *via* the BPG library (v5.9.8)^44^.

### Software availability

iSubGen is freely available as an R package from CRAN at https://cran.r-project.org/web/packages/iSubGen/index.html or from https://github.com/uclahs-cds/public-R-iSubGen.

## Acknowledgements

PCB was supported by a Terry Fox Research Institute New Investigator Award and a CIHR New Investigator Award, and by the NIH/NCI under awards P30CA016042, 1U24CA248265-01 and 1U01CA214194-01. NSF was supported by a CIHR Canadian Graduate Scholarship, a CIHR Michael Smith Foreign Study Supplement, a Medical Biophysics Excellence University of Toronto Fund Scholarship, by the University of Toronto Geoff Lockwood and Kevin Graham Medical Biophysics Graduate Scholarship and by a Prostate Cancer Canada Philip Feldberg Studentship.

## Competing Interest Disclsoure

At the time of publication NSF was an employee of Hoffman-La Roche Limited (Roche Canada). All contributions to the conception, design, analysis and interpretation of results of this project were completed prior to this employment.

## Author Contributions

Initiated the project: NSF, PCB

Coded iSubGen: NSF

Dataset processing and curation: SH, CL

Performed statistical and bioinformatics analyses: NSF

Data interpretation: NSF, PCB

Wrote the first draft of the manuscript: NSF

Supervised research: PCB

Approved the manuscript: all authors

## Supplementary Figure Legends

**Supplementary Figure 1** | **Consensus integrative similarities association with known subgroups**

(A) Association of breast cancer CISs within training (left of each pair) and testing cohorts (right of each pair) with PAM50 subtypes. (B) Testing cohort receiver operator characteristic curves of random forest classifiers predicting each whether the patients are the PAM50 subtype or not that subtype. (C) Gini importance assessment of the CIS contribution to the random forest classifiers. (D) Association of CISs with TCGA cancer types. CISs were independently calculated from two non-overlapping subsets of twelve cancer types. (E) Association of CIS_mRNA-SNV_ with thyroid cancer subtypes. (F) Testing cohort receiver operator characteristic curves of random forest classifiers predicting each cancer type or not that cancer type. (G) Gini importance assessment of the CIS contribution to the random forest classifiers.

**Supplementary Figure 2** | **Integrative subtype generation details**

Schematic details of iSubGen for *n* patients with three data types as an example. (A) The pairwise comparison of data types created a matrix of CISs. Similarity metrics were varied depending on the data type. The patient similarities relative to one patient were compared for each pair of data types using Spearman’s correlations. For example, the column for patient 1 from the CNA similarity matrix was correlated to the column for patient 1 from the methylation similarity matrix and Spearman’s ρ (represented here as a scatterplot showing correlation between the columns) is the integrative correlation. This correlation was repeated with subsampling to create the consensus integrative similarity (CIS). This created a comparison matrix with one row for each patient that we want to classify and one column for each pair of data types. (B) Although different feature reduction approaches can be used, the independent feature reduction step is demonstrated using an autoencoder for each data type. The bottleneck layer from the autoencoder became columns in the independent reduced feature (IRF) output matrix.

**Supplementary Figure 3** | **Pan-cancer iSubGen classification**

(A) Association between CISs and iSubGen-P classifications. Two subsets of TCGA were run independently to create iSubGen-P and iSubGen-Q classifications. (B,D) The number of patients in each TCGA cancer type and iSubGen classification for the first patient cohort (B) and the second patient cohort (D). Colouring represents the number in the group. (C,E) Association between CIS and iSubGen classification. For each data type pair, CIS was compared between that group and the other iSubGen groups using a Wilcoxon rank sum test. The background colouring of each square is the FDR-adjusted p-value from the comparison and the dot size and colouring represent the difference in median CIS for the comparison. (F) Pearson correlation of the median CIS for iSubGen-P and iSubGen-Q groupings.

**Supplementary Figure 4** | **Breast cancer iSubGen using integrative omics features**

(A) Using the iSubGen, the training cohort of 684 breast cancer patients was classified into five subtypes. (B) Overall survival for the iSubGen-B subtypes. P-value is from a log-rank test. (C) Association between iSubGen-B subtypes and cancer grade. P-value is from a χ^2^ test. (D) Association between iSubGen-B subtypes and tumour size. P-value is from a Kruskal-Wallis rank sum test. (E) Comparison of the iSubGen-B subtypes and PAM50 subtypes in the training cohort (top) and testing cohort (bottom). (F) CIS associations with iSubGen-B. (G) Comparison of the iSubGen breast cancer subtypes and IntClust groups from Curtis *et al*. in the training cohort (top) and testing cohort (bottom). Heatmap colouring represents the number of the patients in each overlap and is scaled differently in each cohort.

**Supplementary Figure 5** | **Breast cancer iSubGen using cancer hallmark and pathway mRNA features**

(A) Scaled CIS for nine breast cancer subtypes based on mRNA associated with cancer hallmarks. Rows are patients and columns are pairs of data types which in this case are cancer hallmark mRNAs. (B) Overall survival associations with iSubGen-H subtypes. P-value is from a log-rank test. (C) Association between PAM50 subtypes and iSubGen-H subtypes in the training cohort (top) and testing cohort (bottom). (D) Association between the IntClust subtypes and iSubGen-H subtypes in the training cohort (top) and testing cohort (bottom). (E) Association between the iSubGen-H and iSubGen-B subtypes in the training cohort.

## Supplementary Tables

**Supplementary Table 1** | **iSubGen-B features and subtypes**

**Supplementary Table 2** | **iSubGen-H features and subtypes**

**Supplementary Table 3** | **iSubGen-BH features and subtypes**

**Supplementary Table 4** | **iSubGen-K features and subtypes**

**Supplementary Table 5** | **iSubGen-L features and subtypes**

**Supplementary Table 6** | **Significant associations between CISs and sex**

**Supplementary Table 7** | **iSubGen-P and iSubGen-Q features and subtypes**

