## Supplementary Figures for "iSubGen: Integrative Subtype Generation by Pairwise Similarity Assessment"

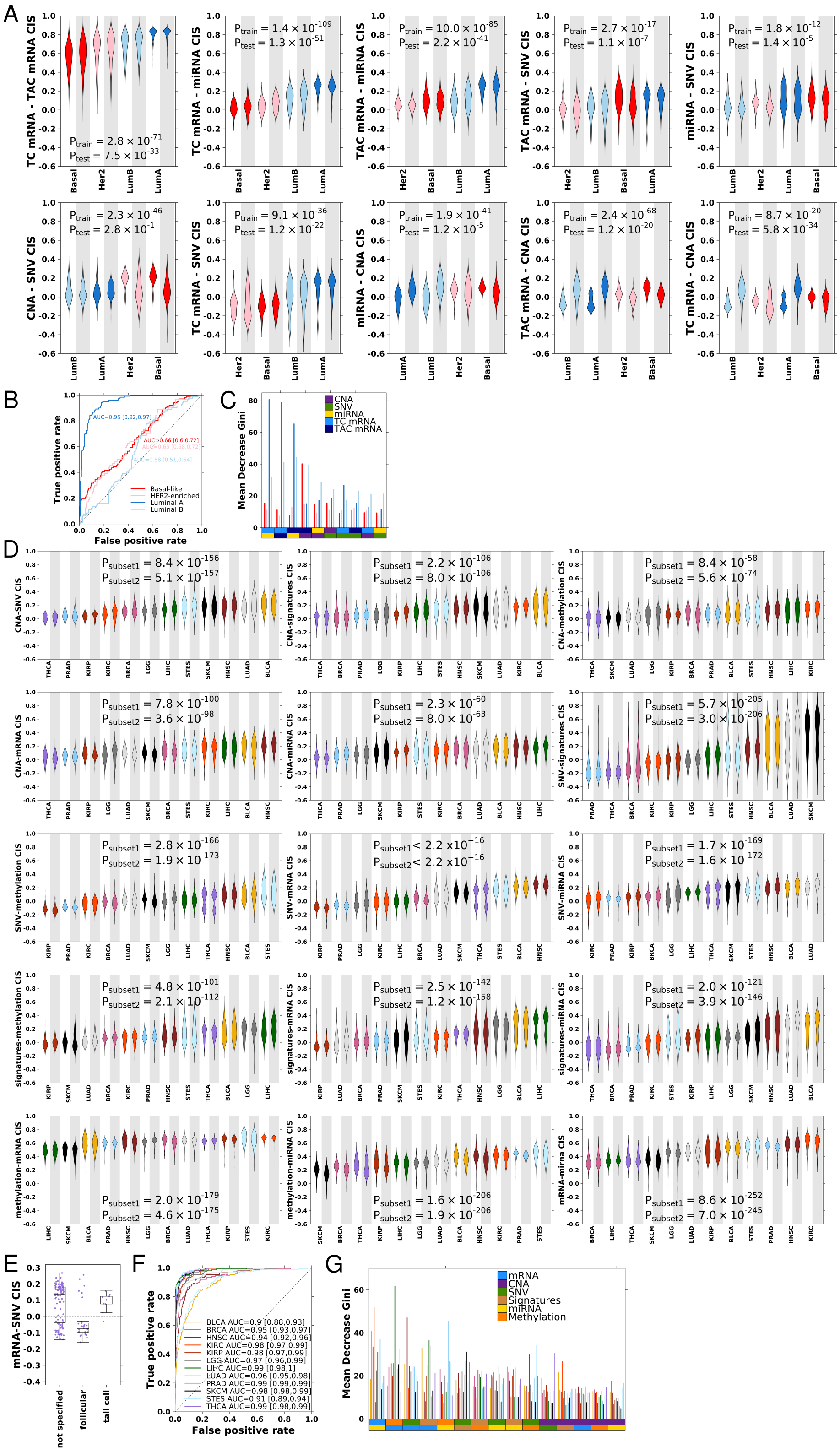

**Supplementary Figure 1 | Consensus integrative similarities association with known subgroups.** (A) Association of breast cancer CISs within training (left of each pair) and testing cohort (right of each pair) with PAM50 subtypes. (B) Testing cohort receiver operator characteristic curves of random forest classifiers predicting each whether the patients are the PAM50 subtype or not that subtype. (C) Gini importance assessment of the CIS contribution to the random forest classifiers. (D) Association of CISs with TCGA cancer types. CISs were independently calculated from two non-overlapping subsets of twelve cancer types. (E) Association of $CIS_{mRNA-SNV}$ with thyroid cancer subtypes. (F) Testing cohort receiver operator characteristic curves of random forest classifiers predicting each cancer type or not that cancer type. (G) Gini importance assessment of the CIS contribution to the random forest classifiers.

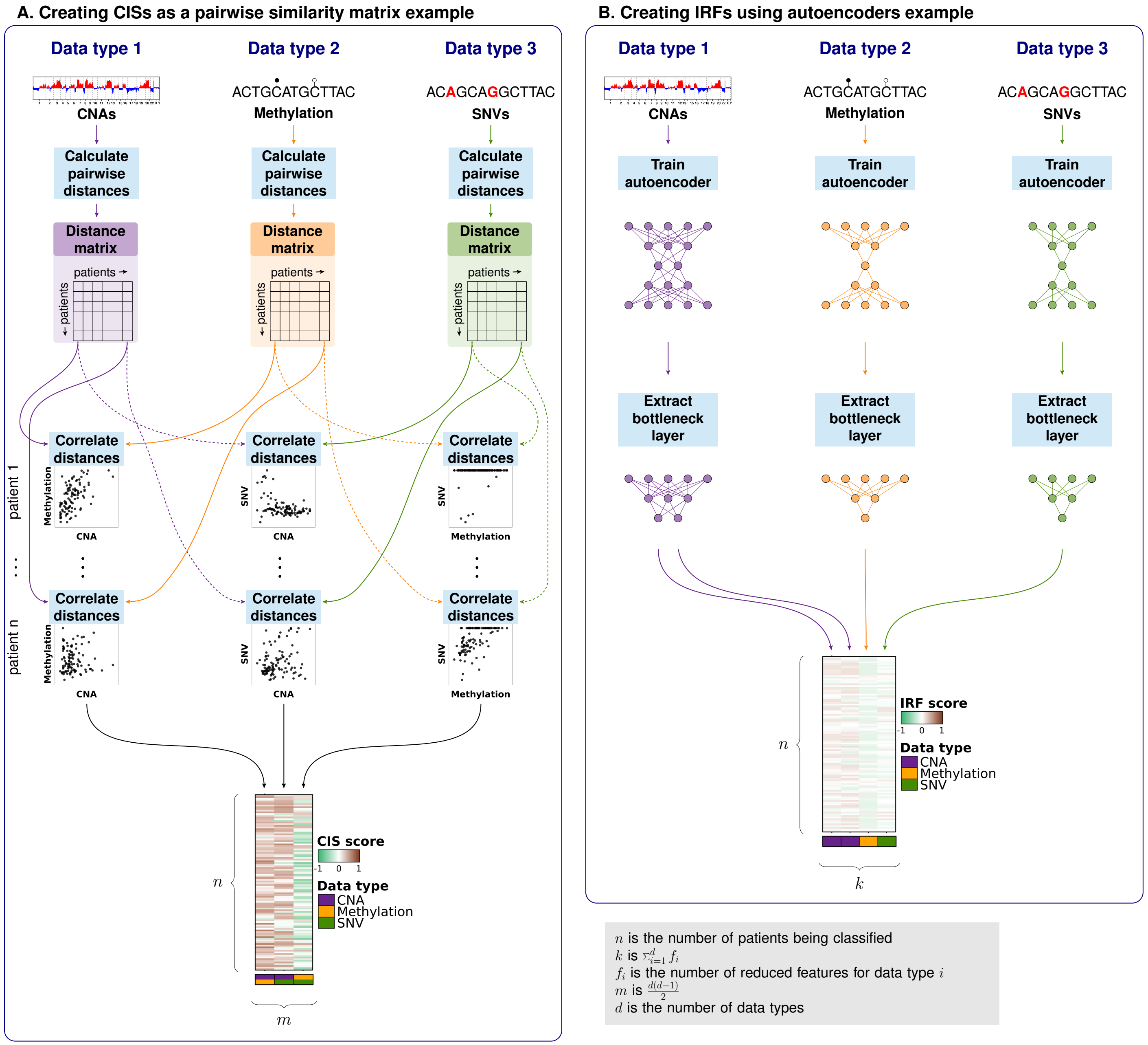


**Supplementary Figure 2 | Integrative subtype generation details.** Schematic details of iSubGen for n patients with three data types as an example. (A) The pairwise comparison of data types created a matrix of CISs. Similaritiy metrics were varied depending on the data type. The patient similarities relative to one patient were compared for each pair of data types using Spearmans correlations. For example, the column for patient 1 from the CNA similarity matrix was correlated to the column for patient 1 from the methylation similarity matrix and Spearmans  (represented here as a scatterplot showing correlation between the columns) was the integrative correlation. This correlation was repeated with subsampling to create the consensus integrative similarity (CIS). This created a comparison matrix with one row for each patient that we want to classify and one column for each pair of data types. (B) Although different feature reduction approaches can be used, the independent feature reduction step is demonstrated using an autoencoder for each data type. The bottleneck layer from the autoencoder became columns in the independent reduced feature (IRF) output matrix.

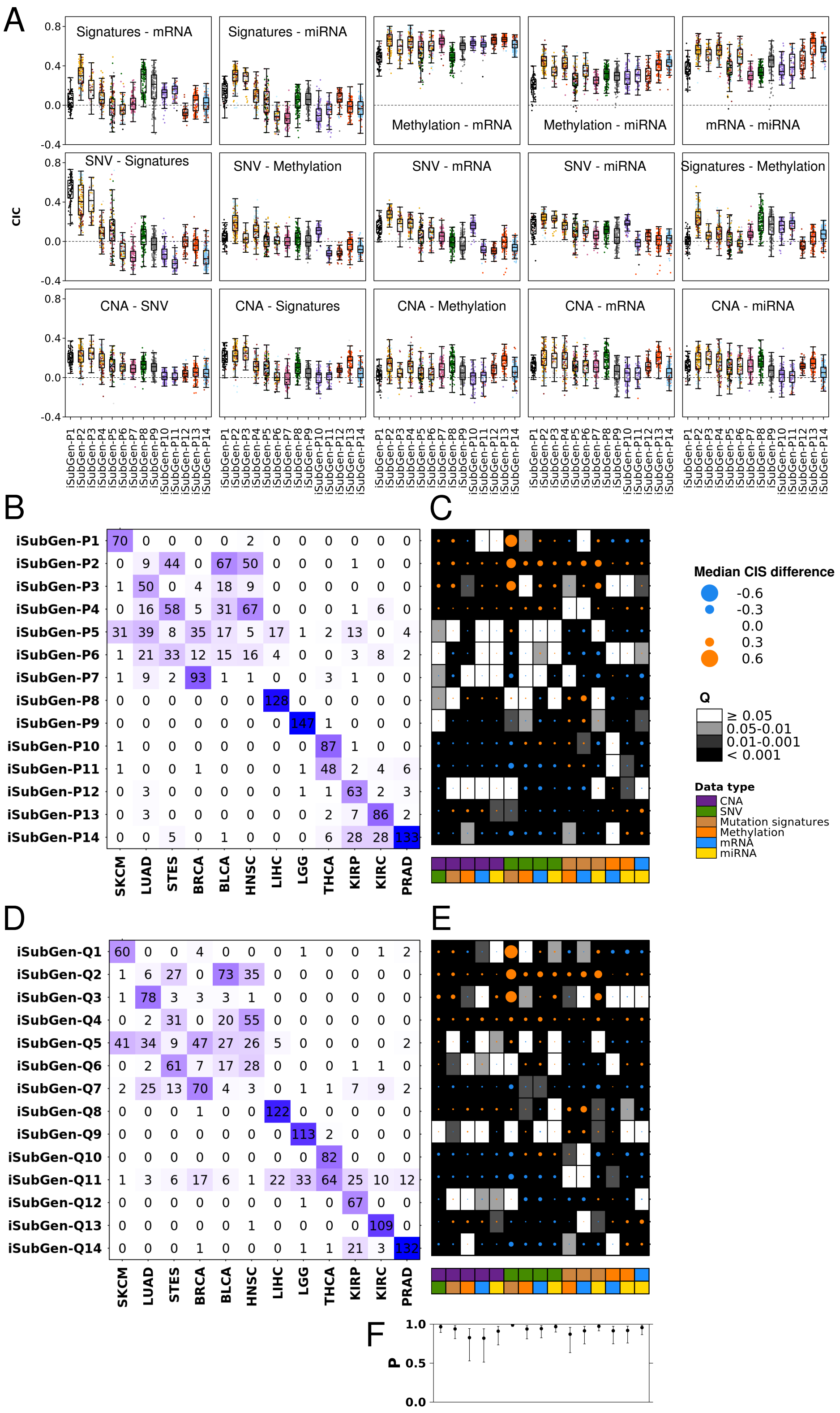


**Supplementary Figure 3 | iSubGen pan-cancer classification** (A) Association between CISs and iSubGen-P classifications. Two subsets of TCGA were run independently to create iSubGen-P and iSubGen-Q classifications. (B,D) The number of patients in each TCGA cancer type and iSubGen classification for the first patient cohort (B) and the second patient cohort (D). Colouring represents the number in the group. (C,E) Association between CIS and iSubGen classification. For each data type pair, CIS was compared between that group and the other iSubGen groups using a Wilcoxon rank sum test. The background colouring of each square is the FDR-adjusted p-value from the comparison and the dot size and colouring represent the difference in median CIS for the comparison. (F) Pearson correlation of the median CIS for iSubGen-P and iSubGen-Q groupings.

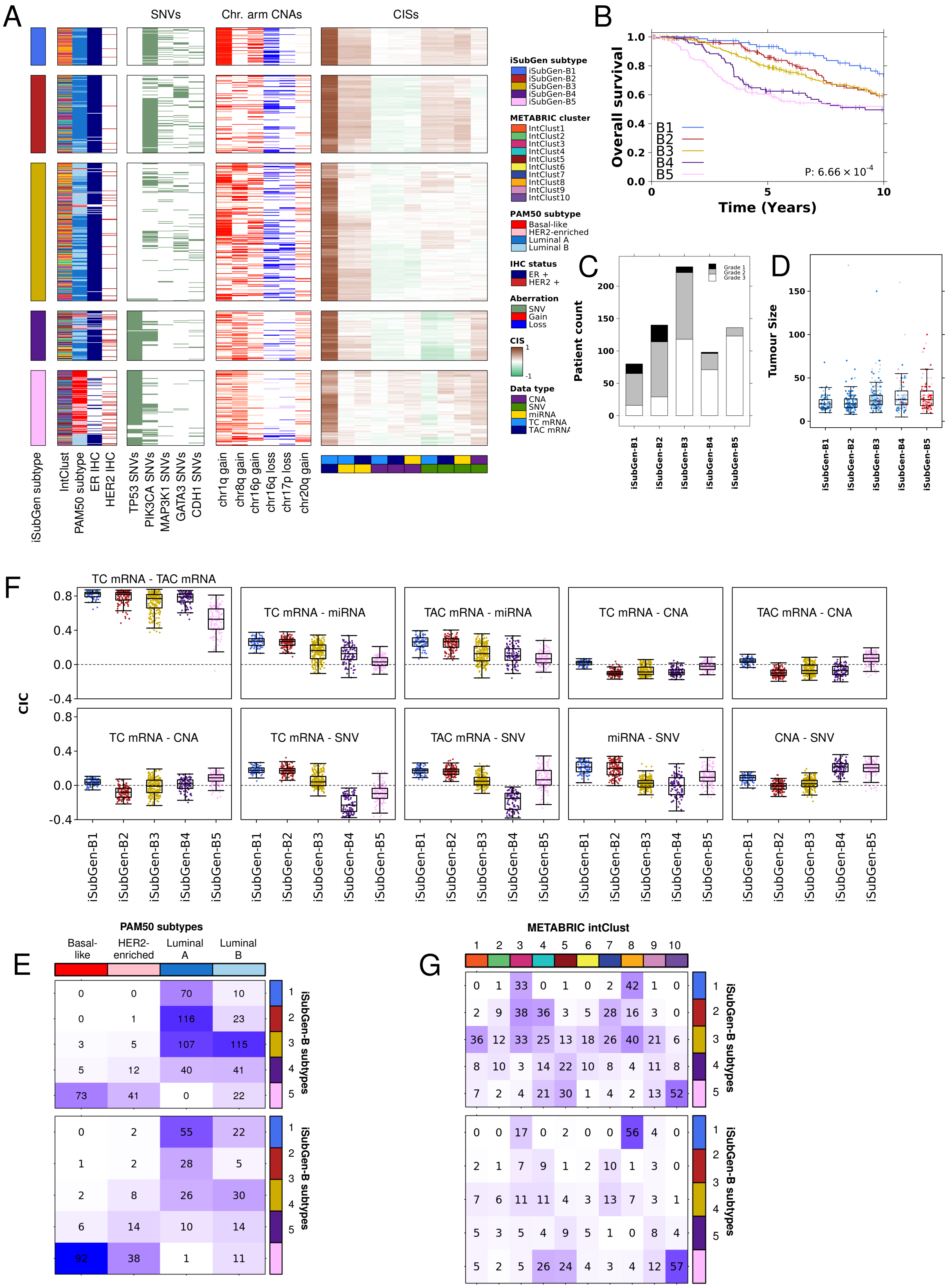


**Supplementary Figure 4 | Breast cancer iSubGen using integrative omics features.** (A) Using the iSubGen, the training cohort of 684 breast cancer patients was classified into five subtypes. (B) Overall survival for the iSubGen-B subtypes. P-value is from a log-rank test. (C) Association between iSubGen-B subtypes and cancer grade. P-value is from a $\chi^2$ test. (D) Association between iSubGen-B subtypes and tumour size. P-value is from a Kruskal-Wallis rank sum test. (E) Comparison of the iSubGen-B subtypes and PAM50 subtypes in the training cohort (top) and testing cohort (bottom). (F) CIS associations with iSubGen-B. (G) Comparison of the iSubGen-B subtypes and IntClust groups from Curtis *et al.* in the training cohort (top) and testing cohort (bottom). Heatmap colouring represents the number of the patients in each overlap and is scaled differently in each cohort.

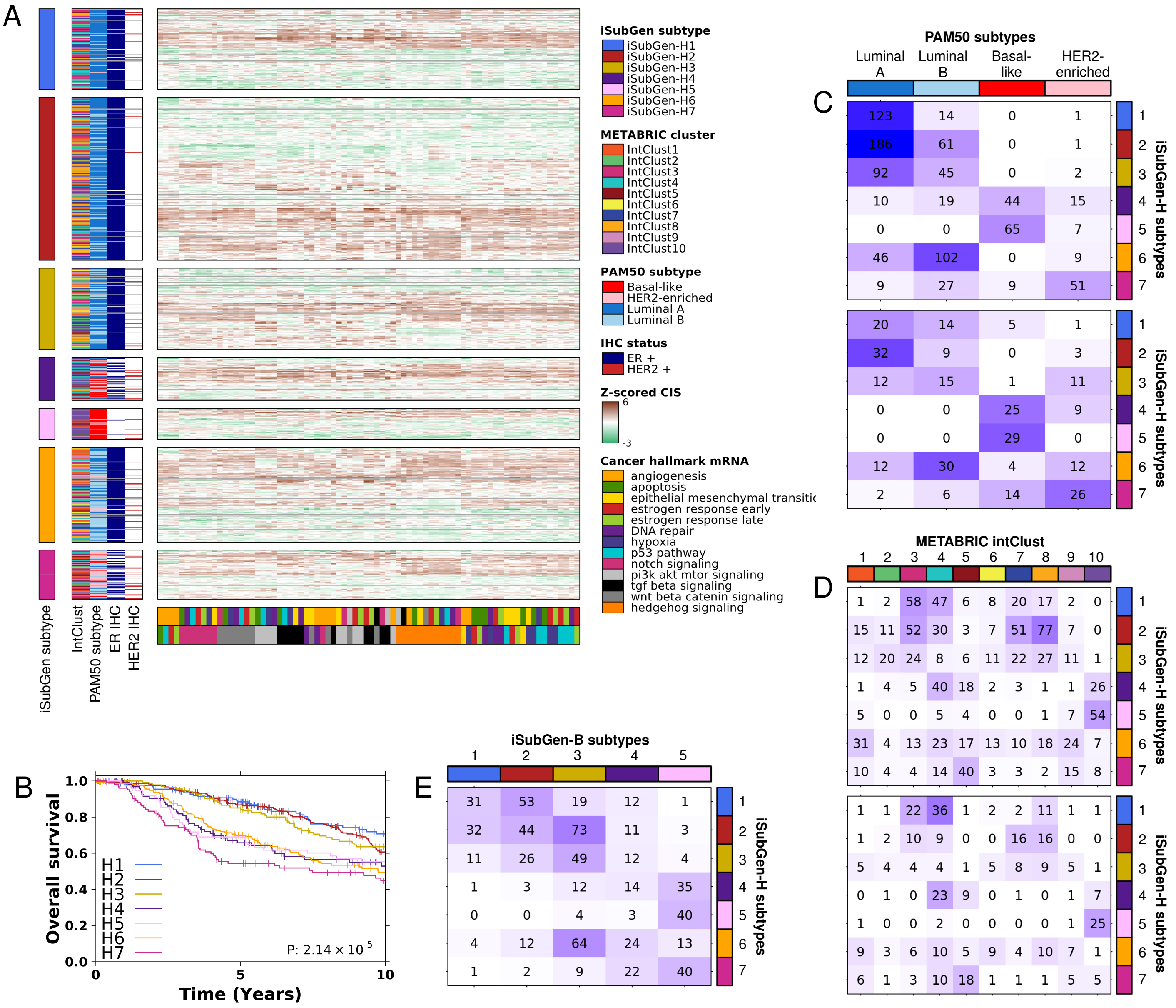


**Supplementary Figure 5 | Breast cancer iSubGen using cancer hallmark and pathway mRNA features.** (A) Scaled CIS for nine breast cancer subtypes based on mRNA associated with cancer hallmarks. Rows are patients and columns are pairs of data types which in this case are cancer hallmark mRNAs. (B) Overall survival associations with iSubGen-H subtypes. P-value is from a log-rank test. (C) Association between PAM50 subtypes and iSubGen-H subtypes in the training cohort (top) and testing cohort (bottom). (D) Association between the IntClust subtypes and iSubGen-H subtypes in the training cohort (top) and testing cohort (bottom). (E) Association between the iSubGen-H and iSubGen-B subtypes in the training cohort.
