## Supplementary Table 6 for "iSubGen: Integrative Subtype Generation by Pairwise Similarity Assessment"

### Supplementary Table 6 | Significant associations between CISs and sex

| **CIS** | **Female**  **Training Cohort**  **Median** | **Male**  **Training Cohort**  **Median** | **Training Cohort**  **Q-value** | **Female**  **Testing Cohort**  **Median** | **Male**  **Testing Cohort**  **Median** | **Testing Cohort Q-value** |
| --- | --- | --- | --- | --- | --- | --- |
| CNA-methylation | 0.05 | 0.11 | 1.2 x 10^-3^ | 0.07 | 0.09 | 0.032 |
| CNA-mRNA | 0.27 | 0.34 | 2.4 x 10^-3^ | 0.27 | 0.31 | 4.9 x 10^-4^ |
| SNV-mRNA | -0.01 | 0.08 | 1.6 x 10^-5^ | 0.00 | 0.05 | 1.0 x 10^-3^ |
| signatures-methylation | 0.01 | 0.07 | 2.9 x 10^-9^ | 0.01 | 0.04 | 1.0 x 10^-3^ |
| signatures-mRNA | 0.10 | 0.18 | 5.5 x 10^-6^ | 0.09 | 0.14 | 4.7 x 10^-4^ |
| methylation-mRNA | 0.58 | 0.49 | 6.2 x 10^-5^ | 0.62 | 0.47 | 2.2 x 10^-11^ |
| methylation-miRNA | 0.25 | 0.23 | 0.023 | 0.25 | 0.22 | 2.8 x 10^-3^ |
